# Machine Learning-Driven Nanopore Sensing for Quantitative, Label-Free miRNA Detection

**DOI:** 10.1101/2025.11.17.688909

**Authors:** Caroline Koch, Seshagiri Sakthimani, Victoria Maria Noakes, Miruna Cretu, David Newman, Richard Gutierrez, Mark Bruce, Julia Gorelik, Nadia Guerra, Joshua B. Edel, Aleksandar P. Ivanov

## Abstract

Nanopore sensors offer exceptional sensitivity for detecting single molecules, making them ideal for early disease diagnostics. In this study, we present a multiplexed nanopore-based assay that combines DNA-barcoded probes with advanced computational analysis to detect microRNAs (miRNAs) with high specificity and quantitative accuracy. Each probe binds selectively to its target biomarker and generates a characteristic delay in the ionic current signal upon translocation through the nanopore, enabling label-free detection.

We evaluated three analytical strategies for classifying delayed versus non-delayed events: (1) moving standard deviation (MSD), (2) spectral entropy (SE), and (3) a convolutional neural network (CNN). While MSD and SE rely on manually defined thresholds and exhibit limited sensitivity, the CNN model, trained on image representations of raw current traces, achieved near-perfect classification performance across all metrics. Grad-CAM visualisation confirmed that the CNN focused on biophysically relevant signal regions, enhancing interpretability and generalisability. All methods produced sigmoidal concentration-response curves consistent with expected binding kinetics, and nanopore-derived delay metrics closely matched RT-qPCR validation data. All three methods were capable of distinguishing between signal classes; however, the CNN model demonstrated superior sensitivity and robustness. This work highlights the importance of data interpretation in nanopore sensing and presents a comparative framework for binary event classification. The findings pave the way for the development of machine learning-driven nanopore diagnostics capable of detecting diverse biomarker types at the single-molecule level.

## INTRODUCTION

Nanopores are single-molecule electrical sensors capable of detecting and characterising individual molecules in real time^1^. This ability to resolve molecular features at the single-molecule level is particularly powerful, as conventional ensemble-based techniques often obscure such detail through averaging effects^2^. Due to their exceptional sensitivity, nanopores have found applications across diverse fields such as environmental monitoring^3^, catalysis studies^4^, DNA-based storage approaches^5,6^, DNA and RNA sequencing experiments^7^ and, most relevant to this study, the detection of biomarkers for distinguishing between healthy and diseased states ^8,9^.

The operational principle involves applying an electrical bias across a nanoscale pore to generate a steady ionic current. When charged analytes translocate through the pore, the ion flow is temporarily disrupted, causing a characteristic change in current. These disruptions encode information about the analyte’s identity, structure, and physicochemical properties^1^.

Due to the complexity of raw ionic current signals in nanopore experiments, a range of computational methods has been developed to extract meaningful information encoded in the data. For biomarker detection, pattern recognition is essential for identifying translocation signatures. Two main challenges must be addressed: specificity and signal interpretation. Since unmodified nanopores lack intrinsic molecular specificity, strategies commonly used include DNA probes^10,11^ or nanopore surface functionalisation^12^ to selectively detect target biomarkers. Signal interpretation typically involves analysis of features such as dwell time and current amplitude^13^, although more complex approaches also examine subpeak structures,^14,15^ and pattern recognition using machine learning (ML)^16^.

In previous work, we developed a nanopore-based biomarker detection strategy using DNA-barcoded probes and nanopore sequencing^17^. This method relies on sequencing a probe’s barcode region and identifying a “delay” in the ionic current signal upon target binding. Events were classified as either delayed or non-delayed, corresponding to the presence or absence of a biomarker. By quantifying the fraction of delayed events, standard curves were generated, allowing for multiplexed detection of microRNAs (miRNAs), proteins, and small molecules. In that study, signal delays were detected using a moving standard deviation (MSD) method; however, this approach relied on manually defined thresholds, limiting both sensitivity and scalability.

This study evaluates three distinct data analysis strategies for interpreting nanopore signals: (1) MSD, (2) spectral entropy (SE), and (3) a convolutional neural network (CNN). While MSD and SE represent traditional, rule-based thresholding techniques, CNN offers a modern, machine learning-based approach that is capable of automating the classification of time-series data. This manuscript explores both paradigms, classical threshold-based methods and advanced deep learning, reflecting the broader shift towards data-driven methodologies in nanopore diagnostics. The performance of all three approaches was benchmarked against a conventional detection technique, reverse transcription quantitative PCR (RT-qPCR). This work introduces a new analytical framework for nanopore signal interpretation and supports the development of ML-driven, high-throughput tools for single-molecule biomarker detection.

## RESULTS AND DISCUSSION

### MSD classification

To demonstrate the biomarker detection concept, previously developed DNA-barcoded probes^17^ were employed to detect hsa-miR-27b-5p, serving as a proof of concept for evaluating different data analysis strategies (Figure 1). These DNA-barcoded probes generate two distinct signal types in nanopore measurements: (1) non-delayed events, indicating the absence of the target miRNA, and (2) delayed events, indicating its presence due to probe-target binding. As an initial analysis, the MSD of the ionic current time series was used to classify events into these two categories. The underlying hypothesis was that miRNA binding to the DNA-barcoded probe would prolong the occupancy of the nanopore by the probe-target complex, leading to reduced signal variability, observable as a decrease in the MSD of the ionic current. In all experiments, the DNA-barcoded probe was incubated with various concentrations of hsa-miR-27b-5p, as described in the Experimental section. The resulting data were used to optimize the MSD method for detecting current delays indicative of target miRNA presence. In brief, the MSD method divides the signal into discrete bins; if the current remains below a predefined threshold across a sufficient number of bins, the event is classified as delayed. The following analyses describe the calibration of these thresholds (Figure 2).

**Figure 1.**
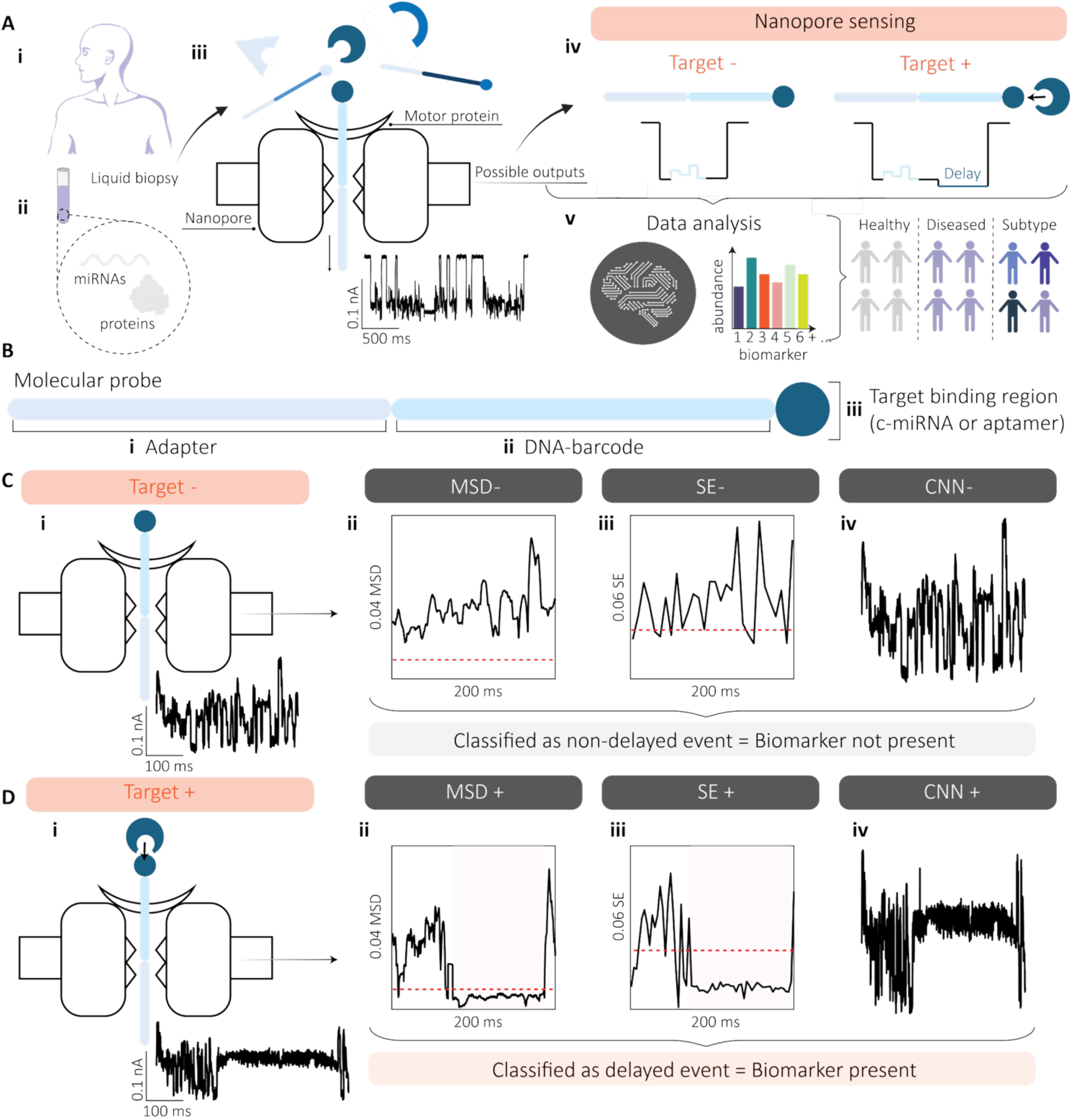
Workflow for biomarker detection using a nanopore assay. **(A)** Schematic overview of the nanopore-based detection process. **(i)** Liquid biopsies collected from patients contain **(ii)** circulating biomarkers such as miRNAs and proteins. **(iii)** These biomarkers are hybridised with DNA-barcoded probes prior to nanopore sequencing. **(iv)** Ionic current traces exhibit no delay in the absence of the target biomarker, but show a characteristic delay when binding occurs. **(v)** Computational analysis of these signals enables quantitative biomarker detection and disease classification. **(B)** Structure of the DNA-barcoded probe, comprising three functional regions: **(i)** an adapter to regulate translocation speed, **(ii)** a unique 35-base barcode sequence, and **(iii)** a target-binding domain, which may be a complementary miRNA sequence or an aptamer for protein recognition. **(C)** Signal classification in the absence of the biomarker. **(i)** The DNA-barcoded probe translocates through the nanopore without delay. To analyse these events, three computational methods were evaluated: **(ii)** MSD, **(iii)** SE, and **(iv)** the CNN model. MSD and SE are rule-based approaches relying on predefined thresholds, whereas the CNN provides automated, data-driven classification of time-series signals. Events that do not fall below the threshold or are classified as class 1 are considered non-delayed. **(D)** Signal classification in the presence of the biomarker. **(i)** Probe-target binding induces a measurable delay during translocation. **(ii)** MSD and **(iii)** SE detect a drop below the threshold, while **(iv)** the CNN classifies the event as class 0. These events are identified as delayed, indicating successful biomarker detection.

**Figure 2.**
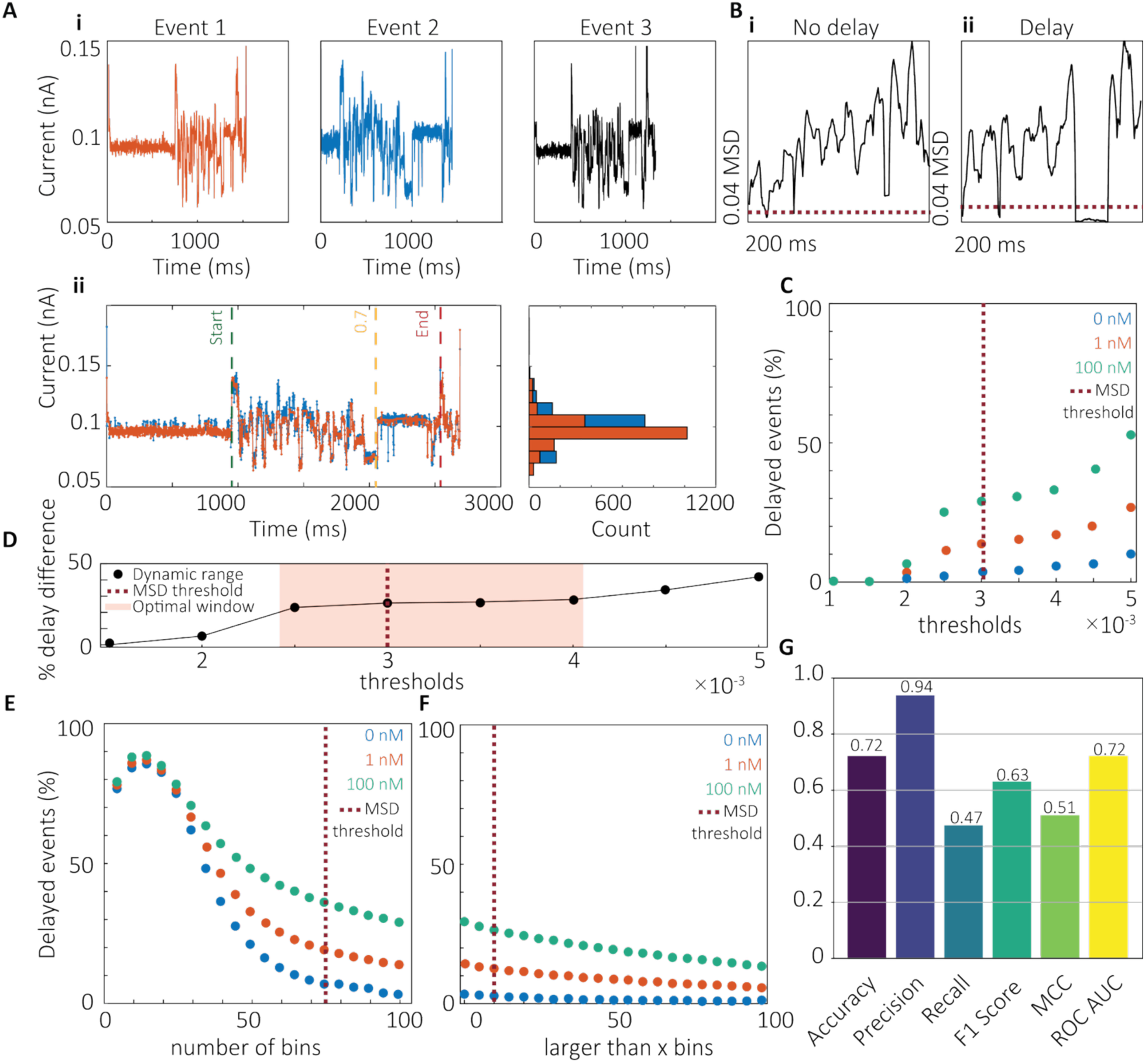
Characterization of the MSD method for miRNA detection using DNA-barcoded probes. **(A)** Delay localisation and reproducibility. **(i)** Representative examples of three randomly selected delayed events. **(ii)** Overlay of two delayed signals showing consistent delay onset at a fractional position of 0.7. **(B)** MSD signal profiles. **(i)** Example of a non-delayed event, where the MSD remains above the threshold. **(ii)** Example of a delayed event, exhibiting a transient drop in MSD below the defined threshold of 0.003. **(C)** Threshold optimisation. Detection thresholds ranging from 0.001 to 0.005 were evaluated across miRNA concentrations of 0, 1, and 100 nM. **(D)** Dynamic range analysis. The difference in delay frequency between 100 nM and 0 nM samples identified an optimal threshold window of 0.0025 - 0.004. **(E)** Bin number effect. Increasing the number of bins from 5 to 100 (with fixed parameters: threshold = 0.003, fractional position = 0.7, minimum delay > 1 bin) revealed an inverse correlation between bin number and delay detection. **(F)** Minimum delay length effect. Varying the minimum number of bins required to classify a delay (0–100 bins), while keeping other parameters constant (threshold = 0.003, fractional position = 0.7, total bins = 100), showed that sensitivity increased with delay length, particularly in miRNA-containing samples. **(G)** Performance metrics of the MSD classifier. Accuracy = 0.72, precision = 0.94, recall = 0.47, F1 score = 0.63, Matthews Correlation Coefficient (MCC) = 0.51, and Receiver Operating Characteristics (ROC) Area under the Curve (AUC) = 0.72, indicating high precision but limited sensitivity due to false negatives.

The fractional position of the delay, which is the delay’s location within the signal expressed as a value between 0 and 1, was first assessed by plotting three representative delayed events to visualise the delay pattern (Figure 2A, i). Dynamic time warping (DTW) was used to quantify signal similarity and locate the delay region (Figure 2A, ii). Delays consistently occurred in the final quarter of the signal, with DTW confirming a strong similarity between events (Euclidean distance = 6.85), which validated a fractional position threshold of 0.7 for delay detection. Representative MSD traces for non-delayed (Figure 2B, i) and delayed (Figure 2B, ii) events showed that only delayed events exhibited a transient drop below the threshold of 0.003 (Supplementary Data 1). To optimise this threshold, datasets from 0, 1, and 100 nM hsa-miR-27b-5p were analysed across a range of thresholds (0.001-0.005). Higher thresholds increased the proportion of delayed events across all concentrations (Figure 2C). The dynamic range, defined as the difference in delay percentage between 100 nM and 0 nM, was plotted to identify the range providing the greatest separation between control and miRNA-containing samples (Figure 2D). The plateau region, corresponding to thresholds between 0.0025 and 0.004, was selected as optimal. A threshold of 0.003 was chosen to maximise discrimination while maintaining a low false-positive rate in controls. Next, the influence of the *number of bins* and the *minimum delay length* (“larger than *x* bins”) was assessed. Varying bin number from 5 to 100 (threshold = 0.003, fractional position = 0.7, minimum delay > 1 bin) revealed an inverse correlation between bin number and delay percentage (Figure 2E). A bin number of 75 was selected, as it produced consistently low delay percentages under control conditions. The *larger than x bins* parameter was then varied (threshold = 0.003, fractional position = 0.7, 100 bins) (Figure 2F). This parameter had little effect on the control data (0 nM) but markedly influenced delay detection in the presence of miRNA. A threshold of >10 bins was selected to balance sensitivity and specificity. To validate the method, an independent dataset of 2,000 manually curated events (delayed and non-delayed) was analysed. The MSD classifier achieved an accuracy of 0.72, precision of 0.94, recall of 0.47, F1 score of 0.63, MCC of 0.51, and ROC AUC of 0.72 (Figure 2G). Further inspection revealed a subset of atypical events exhibiting high-frequency noise within the delayed segment (Supplementary Data 2). These “noisy delays” elevated MSD values, leading to misclassification as non-delayed and contributing to a high false-negative rate. In diagnostic applications, such false negatives are particularly critical, as they may result in missed detections. To mitigate this, an alternative classification approach based on SE was developed to improve the detection of delayed events and enhance overall miRNA identification accuracy.

### SE classification

SE was first introduced by Shannon in 1948 in communication theory and has since been applied to pattern recognition^18^. It can detect potential anomalies or irregularities in time series data by quantifying the level of disorder or unevenness in the frequency distribution and, therefore, was hypothesised to be able to differentiate between non-delayed and delayed events. To achieve this, the same dataset used for MSD calibration was used to optimize SE. This dataset comprised events recorded from DNA-barcoded probes incubated with increasing concentrations of hsa-miR-27b-5p, as described in the Experimental section. The SE method was optimised to detect signal delays indicative of miRNA binding (Figure 3).

**Figure 3.**
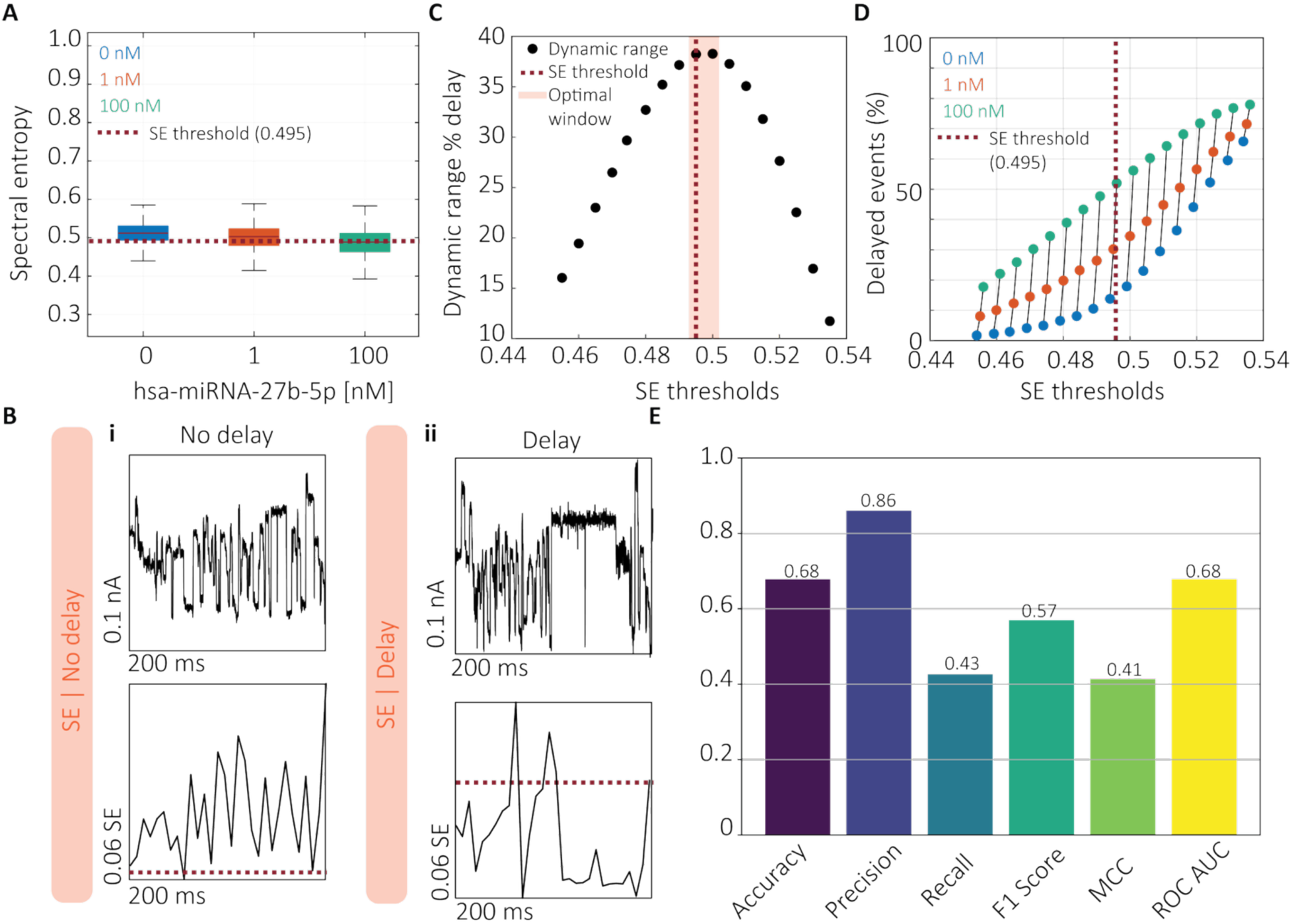
Characterization of the SE method for miRNA detection using DNA-barcoded probes. **(A)** Distribution of SE values for events recorded in the absence (control) and presence of increasing concentrations of hsa-miR-27b-5p (1 nM and 100 nM). The classification threshold for delayed events was set at SE = 0.495, corresponding to the 25th percentile of the control distribution. **(B)** Representative examples of **(i)** a non-delayed event and **(ii)** a delayed event, along with their corresponding SE traces. Delayed events exhibit a distinct drop below the SE threshold, while non-delayed events remain above it. **(C)** Dynamic range analysis (delay at 100 nM minus delay at 0 nM) identified an optimal SE threshold window between 0.495 and 0.505. **(D)** Percentage of delayed events detected across varying SE thresholds for datasets containing 0, 1, and 100 nM miRNA, highlighting the sensitivity of threshold selection. **(E)** Performance metrics of the SE classifier: accuracy = 0.68, precision = 0.86, recall = 0.43, F1 score = 0.57, MCC = 0.41, and ROC AUC = 0.68. These results indicate moderate classification performance, with high precision but limited sensitivity due to a tendency to under-detect true delayed events.

To optimise the SE method’s parameters, the first step was to determine an appropriate threshold for identifying delayed events (Figure 3A). SE were plotted for three datasets corresponding to 0, 1, and 100 nM concentrations of hsa-miR-27b-5p. As anticipated, SE values decreased with increasing miRNA concentration. The 25th percentile of the control (0 nM) distribution, corresponding to an SE value of 0.495, was selected as the classification threshold. Events with SE values below this threshold were classified as delayed, while those above were considered non-delayed. Representative electrical current traces and their corresponding SE profiles for a non-delayed (Figure 3B, i) and a delayed event (Figure 3B, ii) demonstrated that delayed signals exhibited a pronounced drop in SE (Supplementary Data 3). This observation aligns with the theoretical expectations, confirming SE as a reliable indicator of signal delay. Consistent with earlier findings (Figure 2A, ii), miRNA-induced delays were localised to the region preceding the second C3 spacer (fractional position > 0.7). Consequently, SE values from this region were used for classification. The selected threshold (SE = 0.495) was further validated by analysing the dynamic range of delay detection, defined as the difference in delay frequency between 100 nM and 0 nM samples. This analysis identified an optimal threshold window between 0.495 and 0.505 (Figure 3C). Varying the SE threshold across the three datasets revealed that the dynamic range increased up to the selected threshold and declined thereafter (Figure 3D), underscoring the importance of precise threshold selection. Inappropriate thresholds risk capturing barcode-related variability rather than true delay signatures. To evaluate the classifier’s performance, the same dataset of 2,000 events previously analysed with the MSD method was re-evaluated using the SE approach (Figure 3E). The SE classifier achieved an accuracy of 0.68, precision of 0.86, recall of 0.43, F1 score of 0.57, MCC of 0.41, and ROC AUC of 0.68. Despite its utility, the SE method has several limitations. It is sensitive to spectral resolution effects, as the number of frequency components (*N*) used in SE computation strongly influences SE magnitude, complicating cross-dataset comparisons. This effect is particularly pronounced when signal patterns repeat or when sampling resolution is high, leading to length-dependent biases, where shorter signals tend to be classified as non-delayed, while longer ones appear delayed (Supplementary Data 4). A further limitation shared by both the MSD and SE methods is their dependence on manually defined thresholds. Threshold choice substantially affects the estimated proportion of delayed events (Figure 3D), reducing reproducibility and introducing user bias. To overcome this limitation, we developed an alternative, threshold-free classification strategy based on a CNN inspired by the LeNet-5 architecture (Figure 4).

**Figure 4.**
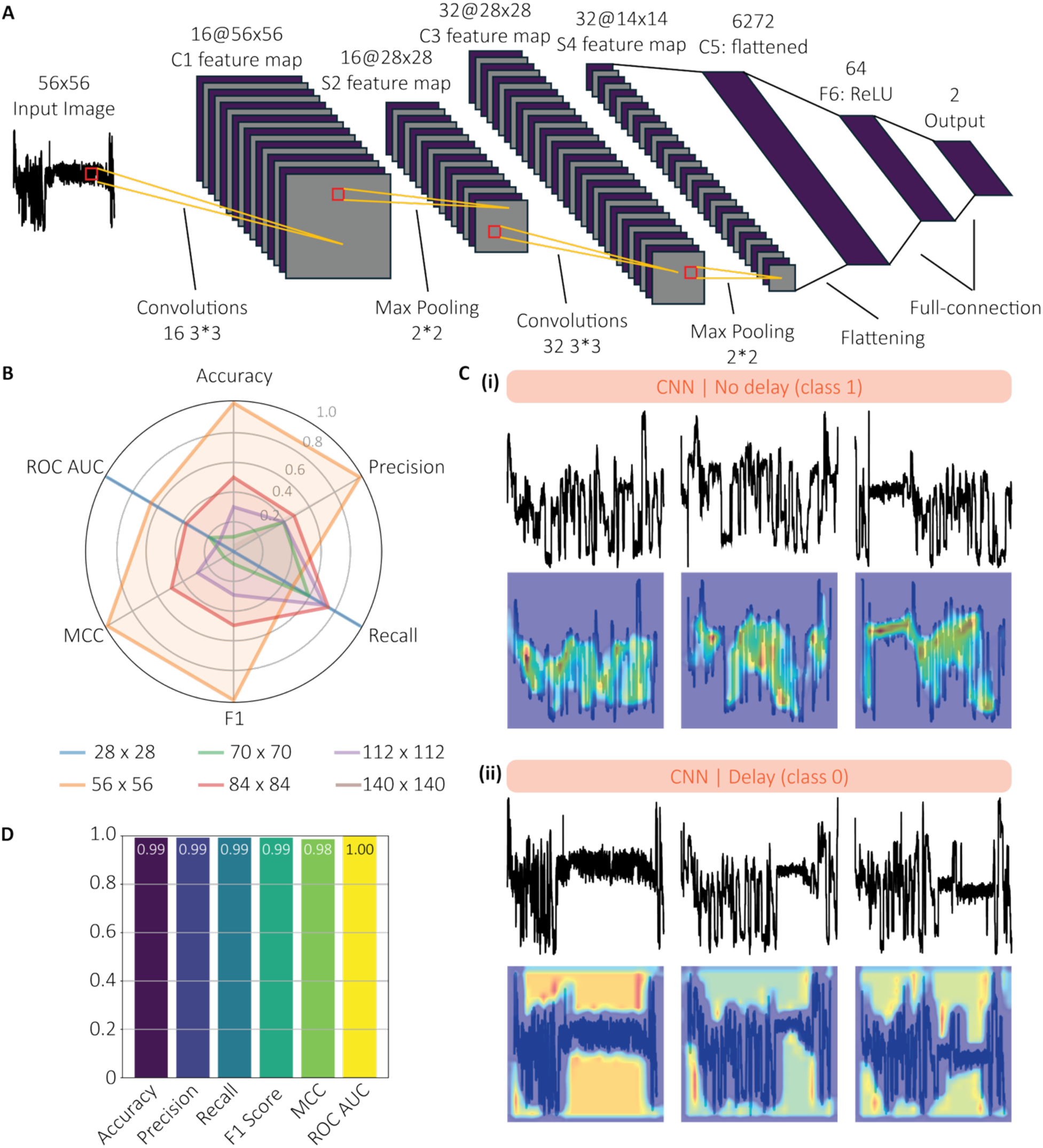
Characterization of the CNN model for miRNA detection using DNA-barcoded probes. **(A)** Architecture of the CNN model, inspired by the LeNet-5 design, comprising two convolutional layers (16 and 32 filters), ReLU activations, max pooling, and a fully connected classification layer. **(B)** Radar plots displaying normalised performance metrics (min–max scaled per metric) for CNN models trained using six different input image resolutions (28 × 28, 56 × 56, 70 × 70, 84 × 84, 112 × 112, and 140 × 140 pixels), highlighting relative performance across resolutions. The 56 × 56 pixel format yielded optimal performance and was selected for subsequent analyses. **(C)** Representative examples of **(i)** non-delayed and **(ii)** delayed events, with corresponding Grad-CAM visualisations. Heatmaps indicate regions of model attention, with delayed events showing strong activation in plateau regions associated with probe-target binding. **(D)** Performance metrics of the CNN classifier: accuracy = 0.99, precision = 0.99, recall = 0.99, F1 score = 0.99, MCC = 0.98, and ROC AUC = 1.00, demonstrating near-perfect classification and superior performance relative to threshold-based methods.

### CNN classification

The CNN model was designed to capture the unique characteristics of signals generated by DNA-barcoded probe translocations, both with and without miRNA binding. The final architecture (Figure 4A) comprised two convolutional layers with 16 and 32 filters, respectively, each followed by ReLU activation and 2 × 2 max pooling for spatial down-sampling. These layers enabled the extraction of local signal features such as amplitude shifts and contour shapes. The resulting feature maps were flattened and passed through a fully connected layer with 64 units before reaching the final classification layer, which outputs one of two classes: delay (class 0) or non-delay (class 1). To optimise input resolution, CNN models were trained using six image sizes (28×28, 56×56, 70×70, 84×84, 112×112, and 140×140 pixels), and performance was evaluated using min–max normalised metrics (Figure 4B). The best performance was achieved at 56×56 pixels, which was therefore selected for subsequent analyses. This architecture was chosen for its simplicity, fast training time, and strong performance on low-resolution input images (Supplementary Data 5). Gradient-weighted Class Activation Mapping (Grad-CAM)^19^ was used to generate saliency maps to identify which regions of the input contributed most to the classification outcome. Grad-CAM was selected over Grad-CAM++^20^ because it produced a smoother, more coherent envelope of the signal, especially for the non-delay representations (Supplementary Data 6). Original current traces were first corrected for signal deviation (Supplementary Data 7) and converted into 56×56 pixel grayscale images (Figure 4C, i-ii). These images were normalized using the mean and standard deviation from model training, then subjected to Grad-CAM analysis to generate heatmaps highlighting areas of model focus (blue = less focus, red = high focus; Figure 4C, i-ii, and Supplementary Data 8). For non-delayed events, the model concentrated on sharp transitions and local fluctuations. For delayed events, attention was focused on extended flat regions in the center of the trace, consistent with unzipping-induced delays. These patterns indicate that the CNN learned to distinguish between delayed and non-delayed signals based on relevant biophysical features, rather than extraneous differences such as barcode sequence. Grad-CAM overlays confirmed that the model’s decisions were grounded in meaningful signal regions, supporting its interpretability and generalizability. To benchmark performance, the same dataset of 2000 events previously analyzed with the MSD and SE methods was re-evaluated using the CNN (Figure 4D). The model achieved an accuracy of 0.99, precision = 0.99, recall = 0.99, F1 score = 0.99, MCC = 0.98, and ROC AUC = 1.00.

Finally, the performance of all three analytical approaches, MSD, SE, and CNN, was compared and benchmarked against RT-qPCR, the current gold standard for miRNA detection (Figure 5).

**Figure 5.**
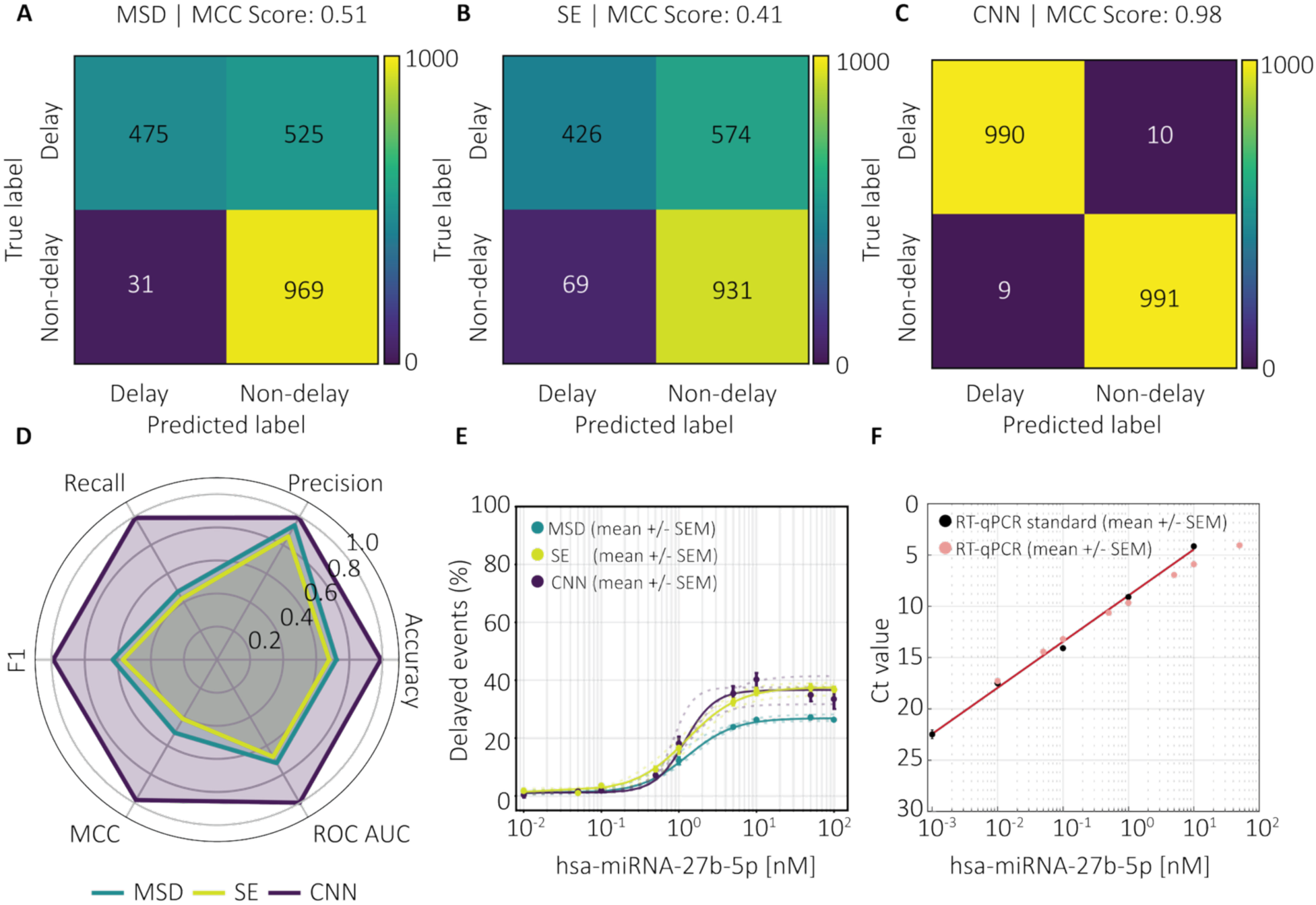
Comparative performance of the MSD, SE, and CNN methods for miRNA detection using DNA-barcoded nanopore probes. (A–C) Confusion matrices illustrating classification outcomes for delay detection using **(A)** the MSD method, **(B)** the SE method, and **(C)** the CNN model. Both MSD and SE exhibit high specificity but reduced sensitivity, whereas the CNN achieves a balanced, accurate classification. **(D)** Radar plots summarising key performance metrics, such as accuracy, precision, recall, F1 score, MCC, and ROC AUC, for all three approaches. The CNN outperforms threshold-based methods across all metrics. **(E)** Concentration–response curves showing the percentage of delayed events as a function of hsa-miR-27b-5p concentration (0–100 nM), fitted using the Hill equation. The MSD method yielded nH=1.43n_H = 1.43 *n*ₕ = 1.43, *K*ₑ = 1.30 nM, *V*ₘₐₓ = 26.88%, *R*² = 0.996; the SE method yielded *n*ₕ = 1.29, *K*ₑ = 1.25 nM, *V*ₘₐₓ = 37.33%, *R*² = 0.998; and the CNN model yielded *n*ₕ = 2.25, *K*ₑ = 1.19 nM, *V*ₘₐₓ = 36.59%, *R*² = 0.965 Data are presented as mean ± SEM (*n* = 3, *n*ₜₒₜₐₗ₋ₑᵥₑₙₜₛ = 1,642,070). All experiments were performed in sequencing buffer. **(F)** RT–qPCR analysis showing the standard curve and the corresponding miRNA concentrations used for nanopore measurements, with decreasing cycle threshold (Ct) values at higher miRNA concentrations.

The confusion matrix showed that the MSD method tends to miss true delays, resulting in low sensitivity but high specificity and precision (Figure 5A). Similarly, the SE method performs well at identifying non-delays but frequently fails to detect true delays, as indicated by its moderate MCC of 0.41 (Figure 5B). These findings suggest that both MSD and SE are prone to false negatives, underscoring a key limitation in sensitivity. In contrast, the CNN model exhibited a markedly improved balance between precision and recall, achieving near-perfect classification accuracy (Figure 5C). This strong performance indicates that the deep learning approach effectively captures complex signal patterns directly from raw traces, enabling more consistent and scalable delay detection than threshold-based methods. To quantitatively compare model performance, standard metrics such as accuracy, precision, recall, F1 score, MCC, and ROC AUC were evaluated (Figure 5D). The CNN model achieved values close to 1.0 across all parameters, demonstrating both exceptional predictive power and balanced classification. By contrast, the MSD and SE methods displayed lower and more variable performance, particularly in recall, confirming their tendency to underdetect true delays. Collectively, these results highlight the superior performance of the CNN model for this classification task. Comparison of the concentration-percentage delay relationships further emphasized the differences among methods (Figure 5E). All three approaches yielded sigmoidal response curves, with percentage delay increasing in a concentration-dependent manner, consistent with expected binding kinetics. The CNN and SE methods demonstrated greater sensitivity at lower miRNA concentrations and earlier saturation than MSD, indicating improved capacity to detect subtle delay signals. The close overlap of the fitted curves and the low standard errors across replicates indicate high reproducibility and robustness of the nanopore-based detection strategy. Finally, RT-qPCR validation confirmed the concentration-dependent detection of hsa-miR-27b-5p (Figure 5F). A standard curve generated from a series of target concentrations (0-50 nM) showed the expected decrease in cycle threshold (Ct) values with increasing miRNA levels. The strong correspondence between nanopore-derived delay metrics and RT-qPCR results supports the accuracy and quantitative reliability of the nanopore assay for label-free miRNA detection.

## CONCLUSION

This study presents a comparison of three approaches for classifying delayed and non-delayed events in nanopore signals generated using DNA-barcoded probes. In nanopore-based detection, a delay in ionic current indicates target binding. Delayed events were identified using the MSD and SE methods, both of which relied on manual thresholding. While effective, both require fine-tuning for each new analyte, limiting their scalability and introducing user bias.

To address these limitations, we developed a lightweight CNN model, inspired by the LeNet-5 architecture, for robust binary classification of nanopore events. The model was trained on 4500 image representations of raw current traces, with 2000 manually curated for blind testing. Pre-processing steps removed extreme outliers and preserved signal dynamics within a 200-700 pA range. Various resolutions were evaluated, with the 56×56 format achieving the best performance across all metrics, including precision, recall, F1 score, AUC-ROC, and MCC. Grad-CAM visualisation confirmed that the CNN focused on plateau regions associated with delays, avoiding barcode-related signal regions. A comparative analysis showed that the CNN consistently outperformed MSD, particularly in terms of sensitivity, validating its superior detection capability. However, the CNN model’s generalisation is currently limited to patterns learned from specific probe configurations, such as delayed events preceding the C3-peak. Future improvements could involve hybrid architectures combining CNNs with time-series models or transformer-based encoders, and real-time implementation for live signal classification.

In conclusion, this study demonstrates that nanopore-based detection, combined with advanced computational analysis, enables robust, quantitative miRNA profiling. Among the three evaluated models, the CNN approach achieved the highest accuracy and sensitivity, outperforming MSD and SE in classifying true delayed events and capturing concentration-dependent signal changes. The strong agreement between nanopore-derived metrics and RT-qPCR validation further supports the method’s quantitative reliability across a wide dynamic range.

## EXPERIMENTAL SECTION

### Nanopore experiments

All nanopore measurements were conducted using the MinION Mk1B device (Oxford Nanopore Technologies, UK) equipped with R10.4.1 flow cells. The device was connected to a computer with a dedicated NVIDIA graphics card. Prior to each experiment, membrane integrity was assessed using MinKNOW software (version 24.11.10 or later, ONT, UK).

### DNA-barcoded probe and miRNA sequence

The DNA-barcoded probe was designed following the protocol described by Koch et al.^17^. Both the probe and its target miRNA (hsa-miRNA-27b-5p) were synthesised by Integrated DNA Technologies (IDT) (Table 1).

**Table 1:**
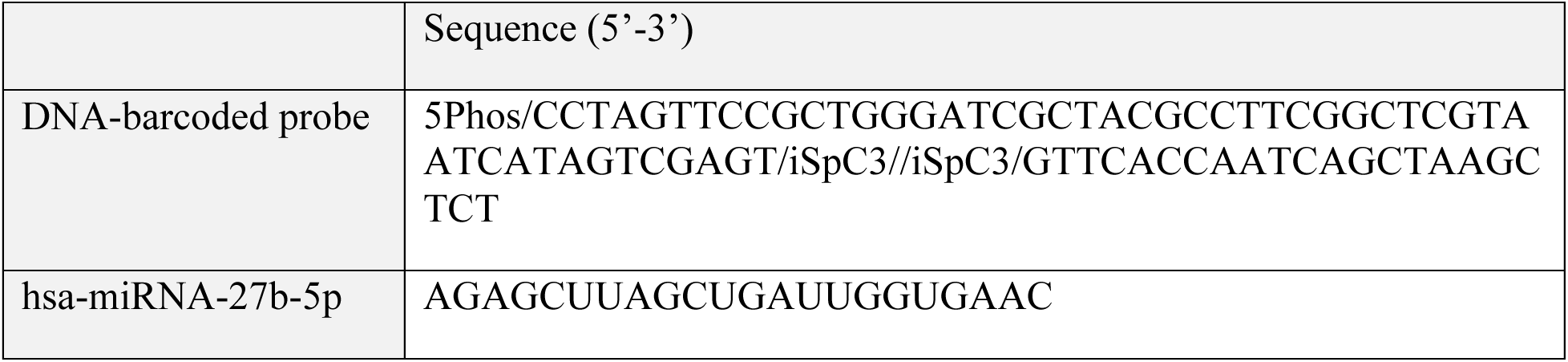
DNA-barcoded probe and hsa-miRNA-27b-5p sequence. 5Phos: Phosphorylation on 5’end, iSpC3: internal C3 spacer.

### Sample preparation

The DNA-barcoded probes (0.225 µL, 20 µM) were incubated with a ligation c-strand (0.6 µL, 25 µM; 5’-CCCAGCGGAACTAGGA-3’) at room temperature (RT) for 1 hour. Subsequently, the mixture was combined with 7.5 µL ligation adapter (ONT, UK) and 8.325 µL TA-ligase (M0367S, New England Biolabs, USA), centrifuged for 1 minute at 3000 rpm, and incubated for 20 minutes at RT on a HulaMixer.

Purification was performed using the solid-phase reversible immobilisation (SPRI) method with Ampure XP beads (A63880, Beckman Coulter, USA). Beads (23.3 µL) were added, followed by two wash steps with 15 µL short fragment buffer (ONT, UK). The beads were then resuspended in 20 µL elution buffer (ONT, UK), releasing the ligated DNA-barcoded probes into solution. Magnetic separation was used to remove the beads, yielding a clear solution of purified, adapter-ligated probes.

### Loading the flow cell

Prior to each run, flow cells were flushed according to the manufacturer’s instructions (ONT, UK). The sequencing mix was prepared in DNA LoBind tubes (Eppendorf, Germany) by combining 37.5 µL sequencing buffer, 12 µL purified DNA-barcoded probes, and 25.5 µL library solution (ONT, UK). For miRNA experiments, the target miRNA was added to the probe mix at concentrations ranging from 0 to 100 nM and incubated for 30 minutes at RT before loading. All experiments were performed in triplicate.

### Nanopore sequencing and basecalling

Data acquisition was carried out using MinKNOW software (version 24.11.10 or later, ONT, UK). Basecalling was performed using the super high-accuracy algorithm provided by ONT.

### Barcode alignment

Basecalled events were aligned to the reference barcode sequence (GGGATCGCTACGCCTTCGGCTCGTAATCATAGTCGAGT) using a local alignment algorithm with scoring parameters: match +5, mismatch -4, and gap -8. Events were further filtered based on the following criteria:

a. Minimum alignment threshold: at least 15 bases matched to the reference
b. Mismatch threshold: no more than 3 mismatches allowed
c. Initial sequence accuracy: alignment must begin with the barcode’s initial “GGG” motif
d. Initial base mismatch filter: a maximum of 3 mismatches permitted within the first 10 bases

Filtered events were subsequently analysed using the classification algorithms to identify signal delays.

### Data analysis

Three methods for detecting the delay were tested, including 1) the MSD of the signal, 2) SE, and 3) a CNN model.

#### 1. MSD

The first approach for detecting a delay in the electrical current signal was based on determining the MSD of the signal. The MSD is defined as:

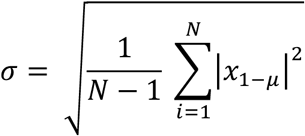

where x is a vector containing N scalar observations and µ is the mean of x. Each standard deviation was calculated over a sliding window of length k across neighbouring elements of xi. The MSD was calculated over k (number of bins), as well as other parameters outlined in Table 2. These parameters were used to classify an event as delayed or non-delayed.

**Table 2:** MSD thresholds. Standard MSD thresholds used in all experiments to determine whether an event was delayed or non-delayed.

|  | Parameter | Threshold |
| --- | --- | --- |
| 1 | $\sigma$ | $< 0.003$ |
| 2 | Total number of bins | 75 |
| 3 | $k$ (bins) | 10 |
| 4 | Delay fractional position | 0.7 |

The parameters indicate 1) a minimum drop of the MSD to 0.003, 2) the total number of bins used for each event, 3) a length condition on the delay pattern, and 4) the fractional position of the delay within the MSD profile.

#### 2. SE

The second approach for detecting a delay in the electrical current signal was based on determining the SE, which quantifies the power distribution within a signal by encoding its spectral attributes. Using the power spectrum of a signal, SE is defined as:

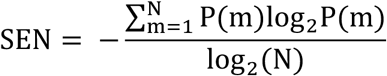

with

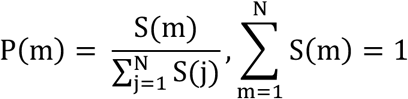

where N is the number of frequency components, S(m) = |X(m)|^2^ and X(m) is the discrete Fourier transformation of the signal x(n). The denominator, log2(N), serves as a normalization factor, ensuring that the SE is constrained within the range [0, 1]. This normalization enables comparability between signals of varying lengths. It represents the maximum theoretical value of SE, achieved when all N frequencies contribute equally to the power spectrum. In the definition of SE provided, P(m) quantifies the percentage contribution of the m^th^ frequency to the spectrum, essentially representing the probability that the signal contains that specific frequency. In the time domain, a flat signal corresponds to a power spectrum with few distinct frequency components, resulting in low SE values. Conversely, SE increases for signals with significant fluctuations in amplitude over time.

To compute SE in MATLAB, the pentropy function was used. This function operates on a vector x sampled at a predefined rate. It first calculates the spectrogram of the input time series and subsequently computes the Shannon entropy. Additionally, it returns instantaneous SEs, providing SE as a function of time. If a time-frequency spectrogram is known, denoted as S(t, f), the probability distribution of frequencies in the power spectrum can be described according to:

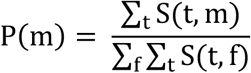

To compute instantaneous SE, the probability distribution at time t becomes:

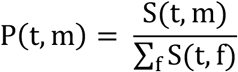

Therefore, the normalised SE at time t reads as:

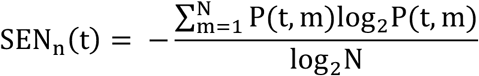

Instantaneous SEs were employed in this manuscript throughout. A threshold of 0.495 was applied: events with SE values dropping below this threshold were classified as delayed.

#### 3. Convolutional Neural Network

The data for training, testing, and validation of the CNN model were selected from sequencing runs using various DNA-barcoded probes with varying miRNA concentrations (0nM, 1nM, 10nM, and 100nM) (Supplementary Data 9). These reads were first aligned to their barcode sequences and then classified as delayed or non-delayed events using the MSD method. The classified events were plotted by class (delayed versus non-delayed). These plots were manually reviewed and selected. A total of 4500 images were curated, representing both classes (delayed versus non-delayed), and were distributed across the training, testing, and validation datasets, as shown in Table 3.

**Table 3:** Distribution of delayed and non-delayed events for training, testing, and validation of the CNN model.

| Class | Training | Testing | Validation |
| --- | --- | --- | --- |
| Delay | 1000 | 1000 | 250 |
| Non-Delay | 1000 | 1000 | 250 |

The classifier architecture was inspired by the LeNet-5 convolutional neural network,^21^ originally designed for handwritten digit recognition. The model was implemented using PyTorch (v2.1.2). Different image resolutions were evaluated to determine the optimal input dimension. Both the test and blind datasets (see section ”data to compare models”) were used to assess model performance. All statistical metrics, including accuracy, precision, recall, F1 score, MCC, and AUC ROC were calculated using scikit-learn (v1.5.2).

### Data to compare models

An additional 2000 images (1000 delayed, 1000 non-delayed) were selected without using the MSD to test the model independently of the labelling pipeline (Supplementary Data 9). This dataset was used to evaluate the viability of MSD and the generalisation performance of the model and was labelled as “Blind data.”

### Statistical metrics

**Recall** measures the model’s ability to correctly identify true positive (TP) events, which were events that were truly delayed. It is defined as:

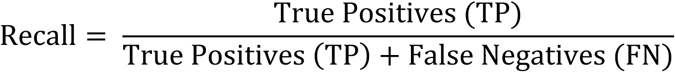

where FN refers to false negatives, or delayed events that were incorrectly classified as non-delayed.

**Precision** measured the correctness of a model’s prediction and is defined as:

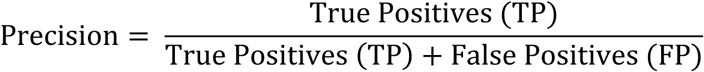

A high precision score indicated that the models produced few FPs and was conservative in assigning the “delay” label.

**F1 Score** is defined as the harmonic mean between precision and recall, which provides a single score that balances both sensitivity and correctness. It is calculated according to:

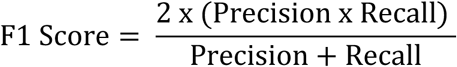

A high F1 score indicated that the model accurately captured true delay events (TP) while also minimizing the false positives (FP).

**MCC** considers all elements of the confusion matrix (TP, TN, FP, and FN) and is defined as:

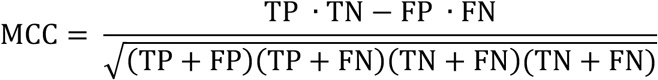

MCC is considered one of the most reliable metrics for evaluating binary classification, especially when the classes are balanced.

**AUC-ROC** assesses the model’s discriminatory power independent of a specific decision threshold. This metric evaluates the true positive rate (TPR) against the false positive rate (FPR) across all thresholds and is defined as:

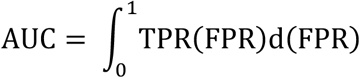

An AUC of 1 indicates perfect separation between classes, while 0.5 indicates random guessing.

**Accuracy** is a common metric and is defined as the ratio of correctly classified instances to the total number of predictions:

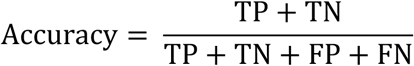

### RT-qPCR

Input RNA was prepared using 1-10 ng of total RNA per 15 µL RT reaction. The components of the RT kit (TaqMan^TM^ MicroRNA Reverse Transcription Kit, 4366596, Thermo Fisher Scientific, USA) were thawed and kept on ice. The RT primer (TaqMan^TM^ Micro RNA Assay, 002174, Thermo Fisher Scientific, USA) was thawed on ice, vortexed briefly, and then centrifuged to collect the contents at the bottom of the tube. The RT reaction mix was prepared according to the manufacturer’s instructions. The RT reaction mix was then centrifuged and placed on ice. Subsequently, 7 µL of RT reaction mix was combined with 5 µL of total RNA in a reaction tube (MicroAmp™ Fast 8-Tube Strip, 0.1 mL, 4358293, ThermoFisher, USA). The tube contents were mixed and centrifuged. Finally, 3 µL of 5×RT Primer (TaqMan^TM^ Micro RNA Assay, 002174, Thermo Fisher Scientific, USA) were added to the reaction tube, centrifuged, and placed on ice before reverse transcription. To perform the RT reaction, the reaction tubes were placed in a PCR machine (StepOnePlusTM, Thermo Fisher Scientific, USA) and incubated using standard cycling conditions, a reaction volume of 15 µL, and the manufacturer’s settings.

To prepare the PCR reaction mix, the PCR Master Mix (TaqMan^TM^ Fast Advanced Master Mix, no UNG, A44359, Thermo Fisher Scientific, USA) was thawed and mixed thoroughly. Then, the PCR reaction mix was prepared according to the manufacturer’s instructions (1 µL TaqMan Small RNA Assay (20x), 10 µL PCR Master mix, and 4 µL nuclease-free water (NFW)). 15 µL of PCR Reaction mix was transferred into each well of an optical reaction plate (MicroAmp® Fast 96-Well Reaction Plate, 4346907, Applied Biosystems, USA). Then, 5 µL of cDNA template or NFW (control) was added to each well. The plate was sealed with optical adhesive film and centrifuged briefly to bring the contents to the bottom of the wells. The manufacturer’s instructions were used to run the RT-qPCR. Experiments were analysed using StepOnePlus software (Thermo Fisher Scientific, USA).

## Acknowledgment

All flow cells were generously provided by Oxford Nanopore Technologies as in-kind contributions.

