## Supplementary Data for "Machine Learning-Driven Nanopore Sensing for Quantitative, Label-Free miRNA Detection"

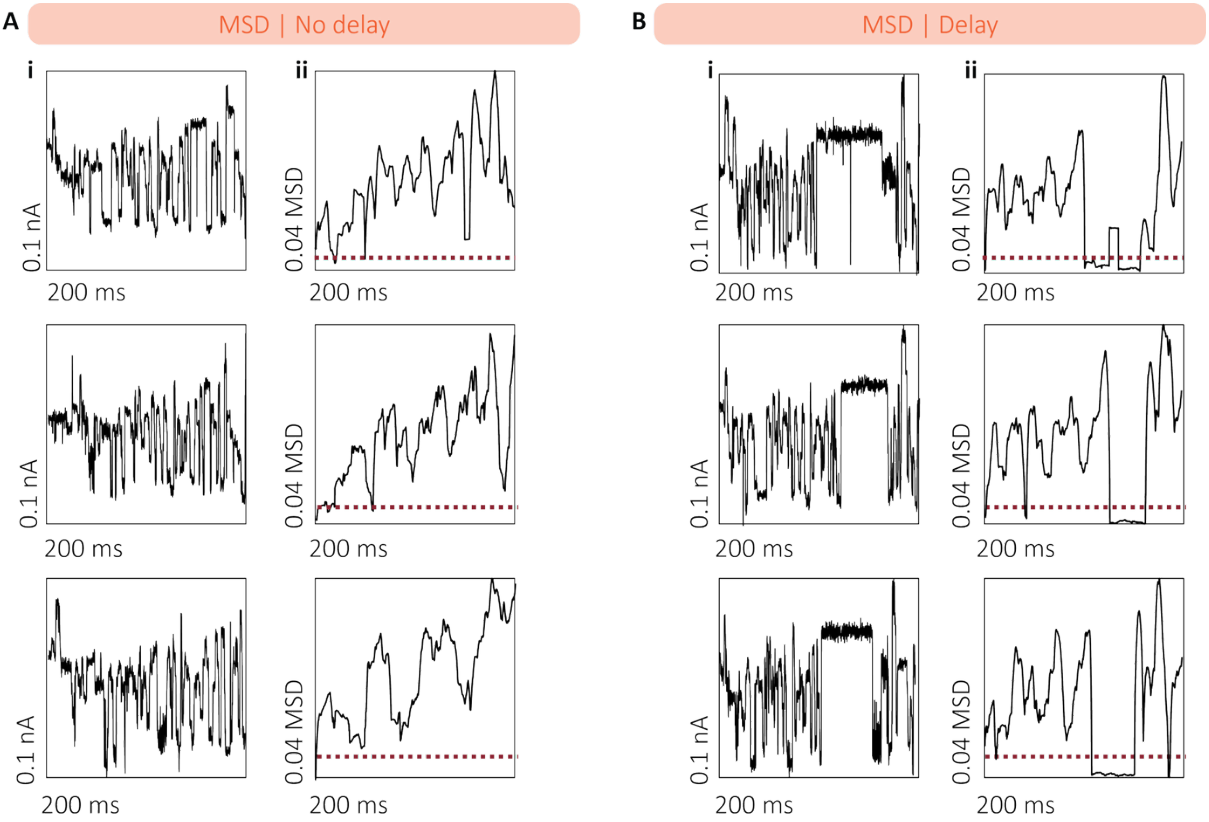


**Supplementary Data 1 | Examples of delayed and non-delayed events using the MSD method. (A)** Representative non-delayed events showing the **(i)** raw current signal and **(ii)** the MSD signal. **(B)** Representative delayed events showing the **(i)** raw current signal and **(ii)** the MSD signal.

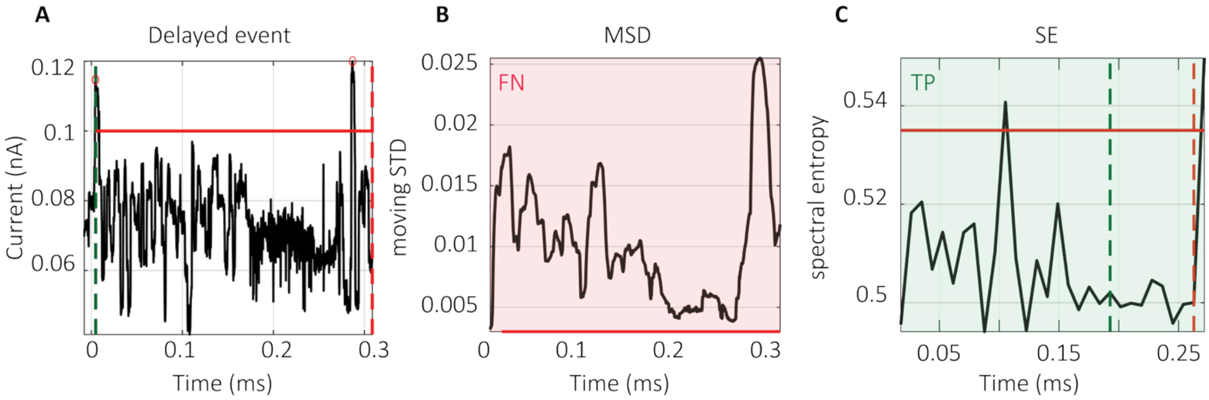


**Supplementary Data 2 | Examples of failed delay detection. (A)** Delayed events exhibiting noisy delay signatures**. (B)** MSD fails to detect the delay due to elevated signal variability. **(C)** SE correctly classifies the same event as delayed, demonstrating improved robustness to noise.

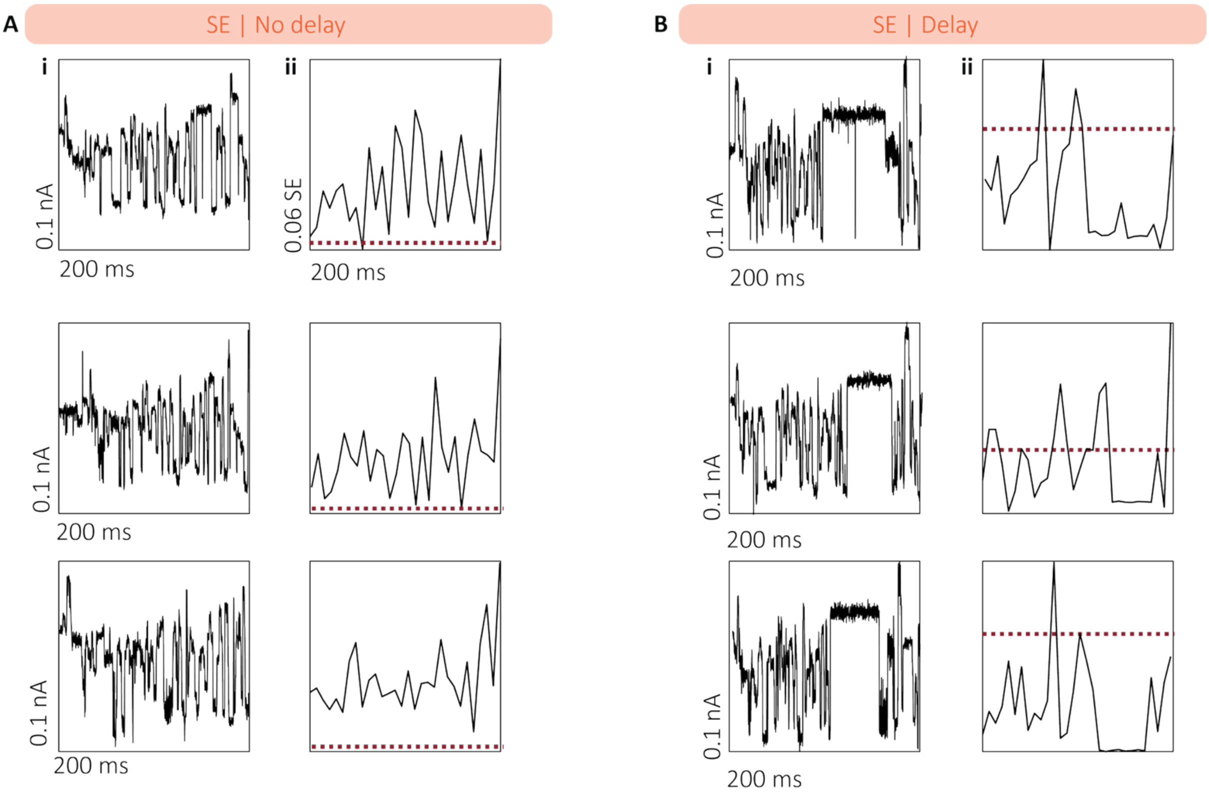


**Supplementary Data 3 | Examples of delayed and non-delayed events using the SE method. (A)** Representative non-delayed events showing the **(i)** raw current signal and **(ii)** the MSD signal. **(B)** Representative delayed events showing the **(i)** raw current signal and **(ii)** the MSD signal.

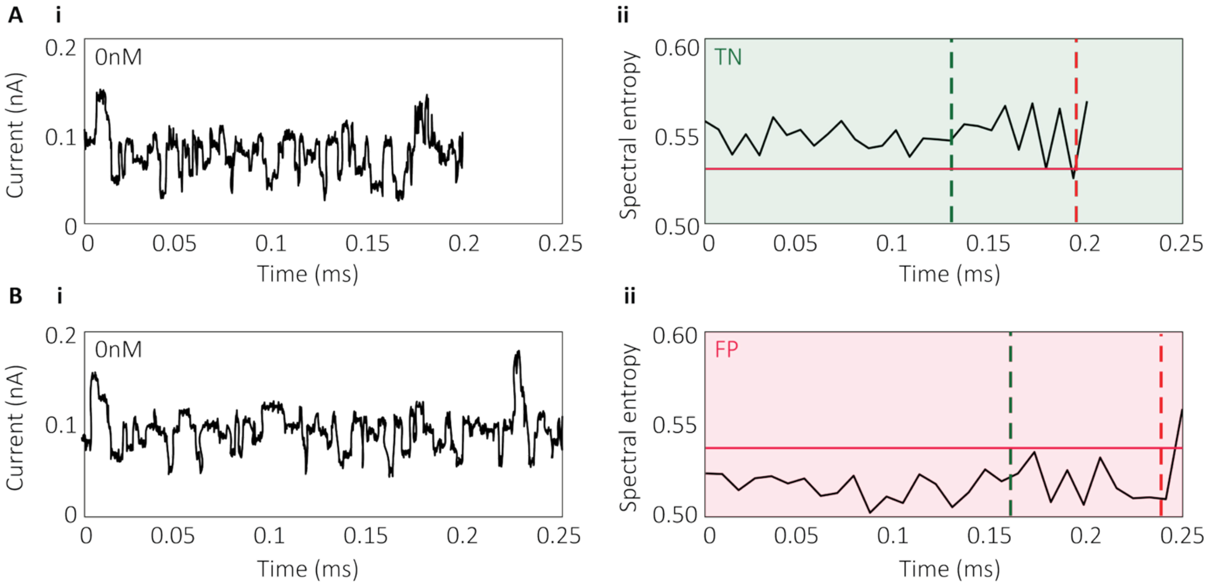


**Supplementary Data 4 | Misclassification of delayed events using SE. (A)** True negative (TN): a non-delayed event correctly classified by SE. **(B)** False positive (FP): a non-delayed event misclassified as delayed due to increased dwell time, highlighting a limitation of SE in distinguishing signal features.

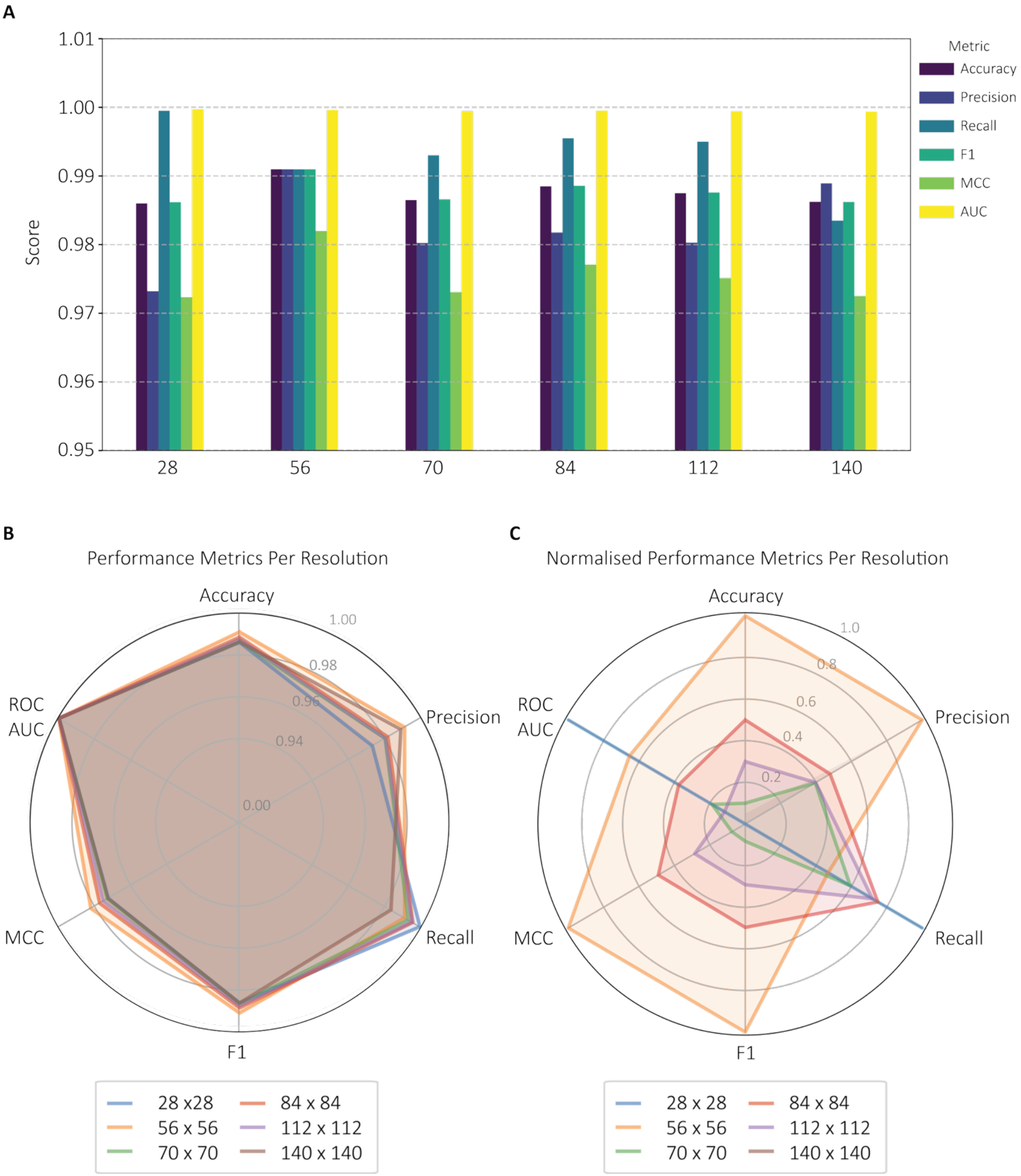


**Supplementary Data 5 | CNN performance across varying image resolutions. (A)** Bar plot comparing raw performance metrics for CNN models trained at six image resolutions 28 × 28, 56 × 56, 70 × 70, 84 × 84, 112 × 112, and 140 × 140 pixels) **(B)** Radar plots of unscaled metric values per resolution. **(C)** Radar plots of normalised metrics (min–max scaling) to highlight relative performance across resolutions.

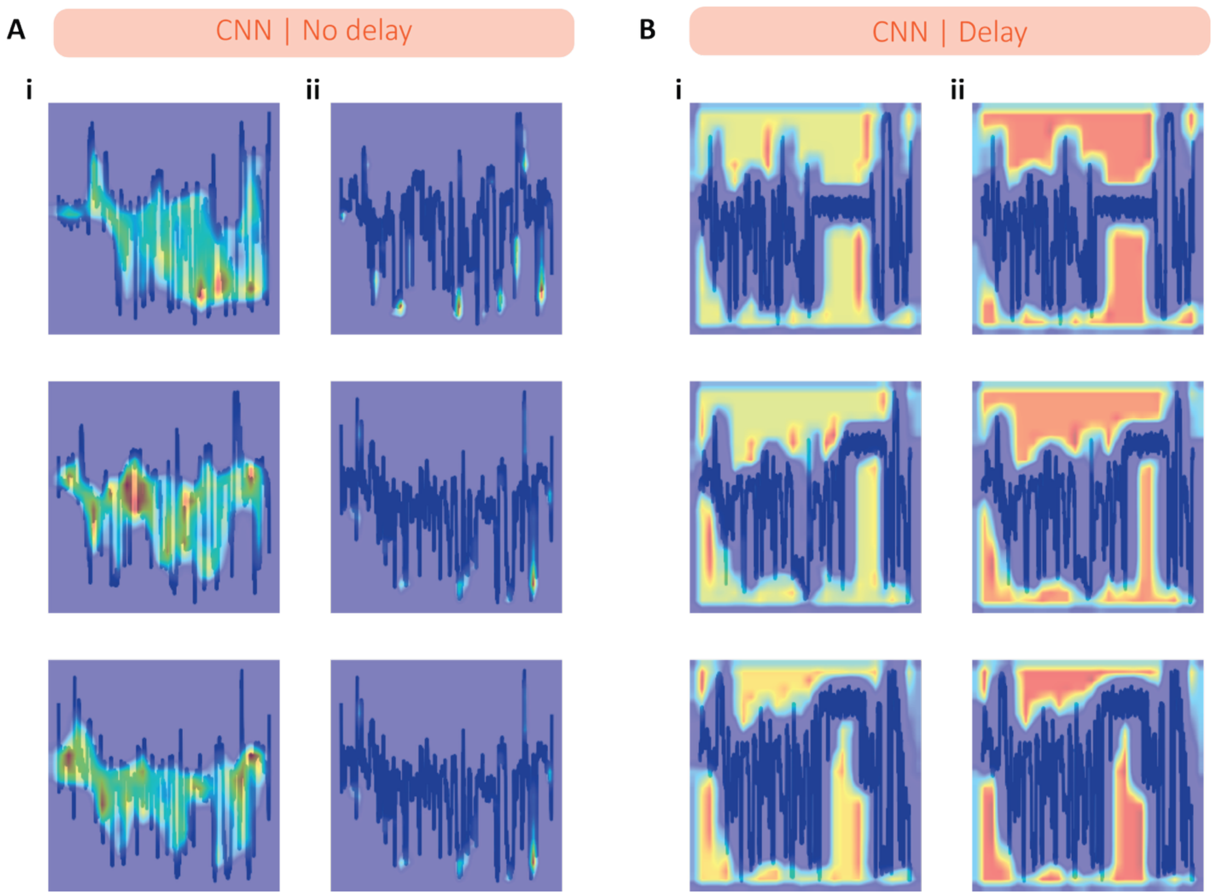


**Supplementary Data 6 | Grad-CAM versus Grad-CAM++ visualisations. (A)** Non-delayed and **(B)** delayed event comparison of saliency maps generated using **(i)** Grad-CAM and **(ii)** Grad-CAM++ for non-delayed events. Grad-CAM was selected for its ability to consistently highlight the full signal envelope, resulting in clearer, more coherent visualisations across both delayed and non-delayed conditions.

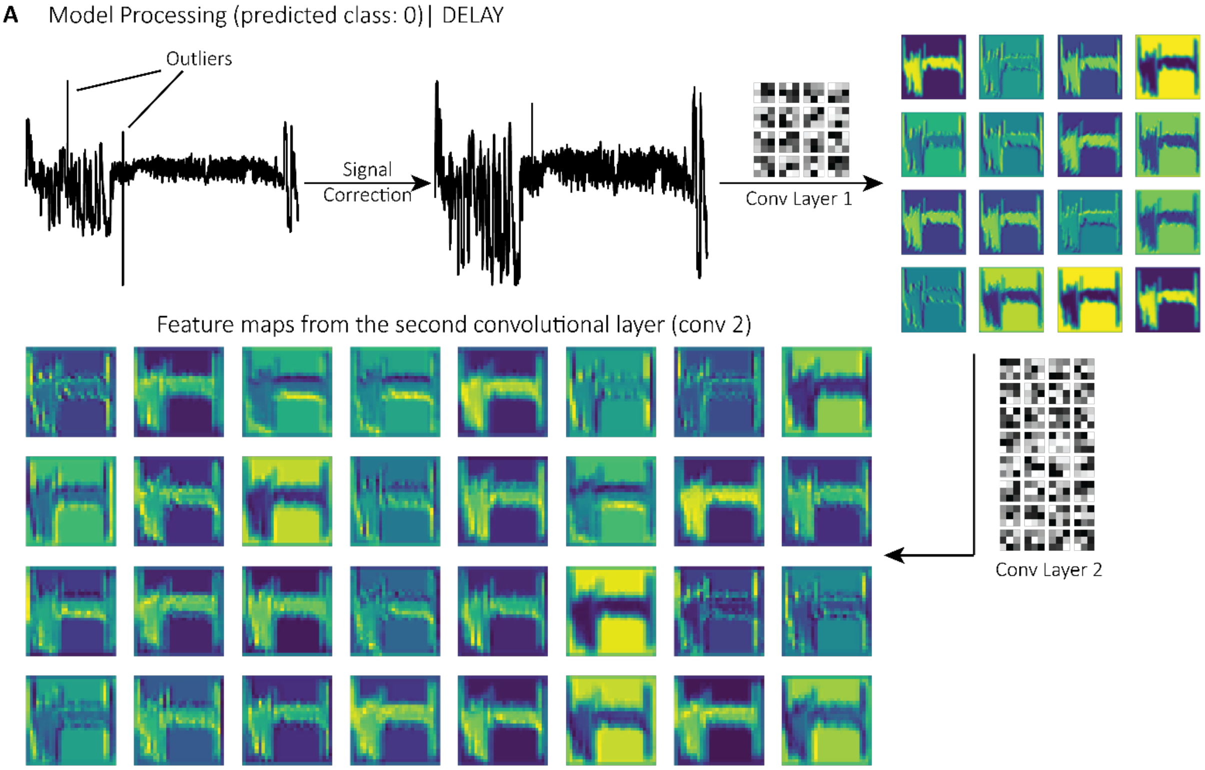


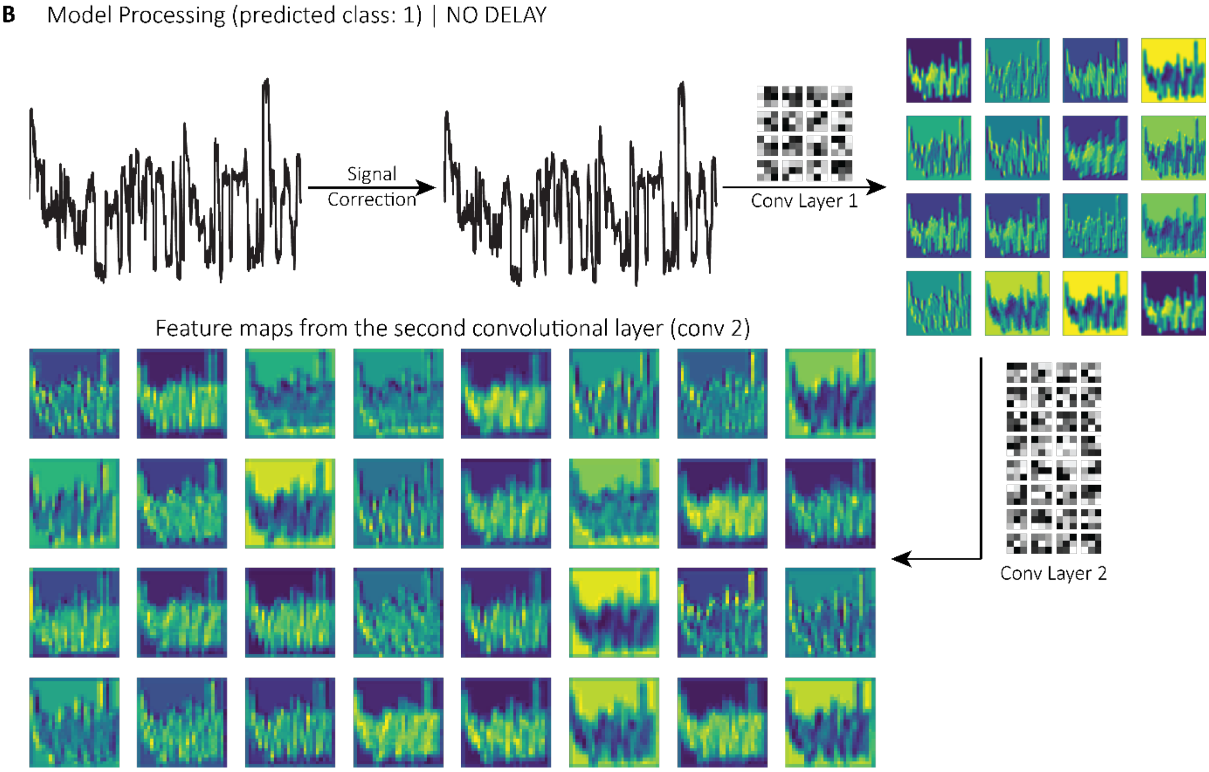


**Supplementary Data 7 | CNN signal processing workflow.** Illustration of the CNN pipeline, showing image transformation through successive convolutional layers. **(A)** Example of a delayed event containing outliers, processed through the signal correction algorithm. **(B)** Example of a clean non-delayed event. The image structure becomes abstracted after the second convolutional layer and is therefore not shown.

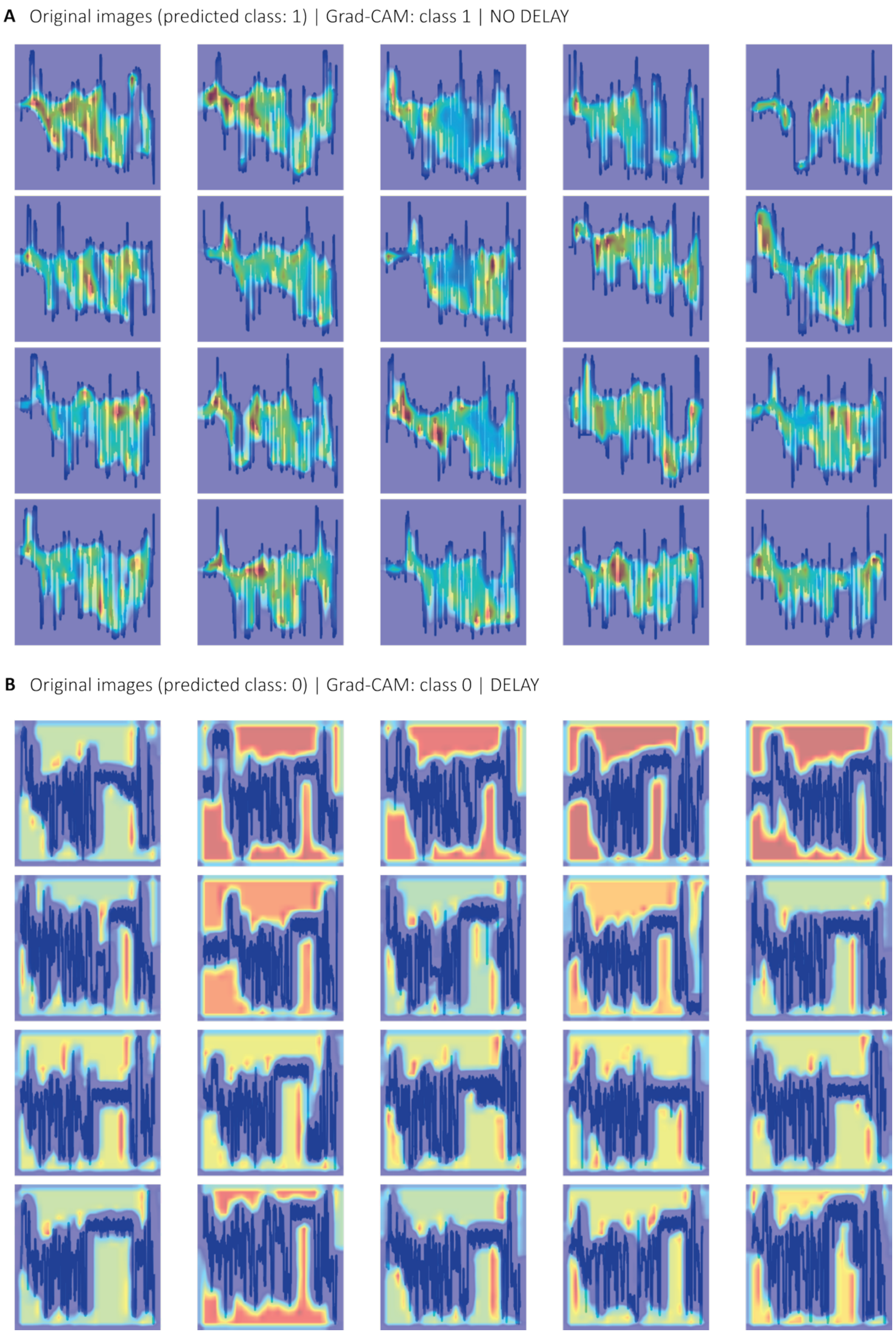


**Supplementary Data 8 | Grad-CAM analysis of CNN attention across multiple events (A)** Saliency maps for 20 non-delayed events. **(B)** Saliency maps for 20 delayed events. The CNN consistently focuses on signal regions associated with delay, confirming the model's interpretability and robustness across diverse examples.

**Supplementary Data 9 | DNA-barcoded probe sequences and miRNA targets used for CNN training and testing.** Table listing all DNA-barcoded probes and their corresponding miRNA targets. Probes and targets highlighted in blue and marked with an asterisk (*) were included in the training dataset.

| DNA-barcoded probe sequence | miRNA target | miRNA sequence |
|---|---|---|
| /5Phos/CCT AGT TCC GCT GGG ATC GCT ACG CCT TCG GCT CGT AAT CAT AGT CGA GT/iSpC3//iSpC3/G TTC ACC AAT CAG CTA AGC TCT | hsa-miR-27b-5p* | rArGrA rGrCrU rUrArG rCrUrG rArUrU rGrGrU rGrArA rC |
| /5Phos/CCT AGT TCC GCT GGG CTA GTG CGC AGT TGT CTC GGC GGA GTT GAG ACT GA/iSpC3//iSpC3/A CCC ACC GAC AGC AAT GAA TGT T | hsa-miR-181b-5p* | rUrArG rCrUrU rArUrC rArGrA rCrUrG rArUrG rUrUrG rA |
| /5Phos/CCT AGT TCC GCT GGG GTT CAC ATC AAG GTC ATA CCG CGA GTT CTA TTT TA/iSpC3//iSpC3/C AGT GTG CGG TGG GCA GGG GCT | hsa-miR-210-5p* | rArCrC rUrGrG rCrArU rArCrA rArUrG rUrArG rArUrU rU |
| /5Phos/CCT AGT TCC GCT GGG GCT TGG GGG ATA GAT GTG CCC CGC GCA TCG GAC CT/iSpC3//iSpC3/C GCG TAC CAA AAG TAA TAA TG | hsa-miR-126a-5p* | rUrGrU rArArA rCrArU rCrCrC rCrGrA rCrUrG rGrArA rG |
| /5Phos/CCT AGT TCC GCT GGG AAC CTT AGG GGC CTC GAA TCT TTG AGA CGA CTA GG/iSpC3//iSpC3/C TAT CTG CAC TAG ATG CAC CTT A | hsa-miR-18b-5p* | rArGrC rCrCrC rUrGrC rCrCrA rCrCrG rCrArC rArCrU rG |
| /5Phos/CCT AGT TCC GCT GGG ATG ACA CAC GTT TTC GAT AGG GAC GCC GAC TTT AA/iSpC3//iSpC3/T GGA CGT TTG CAG GGG AGG TGG | hsa-miR-1306-5p | rUrGrU rArArA rCrArU rCrCrU rArCrA rCrUrC rUrCrA rGrC |
| /5Phos/CCT AGT TCC GCT GGG TGA TAA TAA GAC CTG ACA GAC AAT AGG GAG AAC TC/iSpC3//iSpC3/C GCG TAC CAA AAG TAA TAA TG | hsa-miR-126-5p | rArArC rArUrU rCrArU rUrGrC rUrGrU rCrGrG rUrGrG rGrU |
| /5Phos/CCT AGT TCC GCT GGG ATT AGC GGA ACC AAA CCC AGG AAG GCT TGA AGG CG/iSpC3//iSpC3/A TAC ATA CTT CTT TAC ATT CCA | hsa-miR-1-3p | rUrArG rCrArC rCrArU rCrUrG rArArA rUrCrG rGrUrU rA |
| /5Phos/CCT AGT TCC GCT GGG TAA TTA CTG CCC CAC CAT GAC ATT TTA ATA GCA GT/iSpC3//iSpC3/C TAA CTG CAC TAG ATG CAC CTT A | hsa-miR-301a-5p | rCrArU rUrArU rUrArC rUrUrU rUrGrG rUrArC rGrCrG |
